# The role of maternal effects on offspring performance in familiar and novel environments

**DOI:** 10.1101/2020.01.21.913822

**Authors:** Milan Vrtílek, Pierre J. C. Chuard, Maider Iglesias-Carrasco, Michael D. Jennions, Megan L. Head

**Affiliations:** The Czech Academy of Sciences, Institute of Vertebrate Biology, Brno, Czech Republic; Department of Biological Sciences, Bishop’s University, Sherbrooke, Canada; Research School of Biology, The Australian National University, Canberra, Australia

**Keywords:** genetic correlation, host adaptation, maternal effects, seed beetle, life history

## Abstract

Maternal effects are an important evolutionary force that may either facilitate adaptation to a new environment or buffer against unfavourable conditions. The degree of variation in traits expressed by siblings from different mothers is frequently sensitive to environmental conditions. This could generate a Maternal-by-Environment interaction (M×E) that could inflate estimates of Genotype-by-Environment effects (G×E). We aimed to test for environment-specific maternal effects (M×E) using a paternal full-sib/half-sib breeding design in the seed beetle *Callosobruchus maculatus,* where we split and reared offspring from the same mother on two different bean host types – original and novel. Our quantitative genetic analysis indicated that maternal effects were very small on both host types for all the measured life-history traits. There was also little evidence that maternal oviposition preference for a particular host type predicted her offspring’s performance on that host. Further, additive genetic variance for most traits was relatively high on both hosts. While there was higher heritability for offspring reared in the novel host, there was no evidence for G×Es, and most cross-host genetic correlations were positive. This suggests that offspring from the same family ranked similarly for performance on both host types. Our results point to a genetic basis of host adaptation in the seed beetle, rather than maternal effects. Even so, we encourage researchers to test for potential M×Es because, due to a lack of testing, it remains unclear how often they arise.

## Introduction

Maternal effects modify phenotypes that undergo selection and are therefore a potentially important evolutionary force (Mousseau and Fox, 1998; Räsänen and Kruuk, 2007; Moore *et al*., 2019). They may either facilitate adaptation to novel environments (Fox and Savalli, 2000; Leftwich *et al*., 2019), or buffer against changing conditions through trans-generational phenotypic plasticity (Shama *et al*., 2014). Maternal effects have also been shown to have a range of other implications, such as moderating population dynamics (Plaistow and Benton, 2009), and increasing niche breadth (Van Asch *et al*., 2010). Maternal effects, which can arise due to both genetic and environmental variation among mothers, can be defined as ‘the causal influence of the maternal genotype or phenotype on the offspring phenotype’ (Wolf and Wade, 2009). Genetic maternal effects might increase total heritability and facilitate evolution (Wilson *et al*., 2005; Räsänen and Kruuk, 2007), as opposed to phenotypic (environmental) maternal effects. Teasing these two sources of maternal effects apart is often challenging, however (Kruuk and Hadfield, 2007). Irrespective of their genetic basis, maternal effects increase the degree of similarity between siblings and may thus bias estimates of direct genetic effects upward if ignored (Räsänen and Kruuk, 2007; McAdam *et al*., 2014). This can lead to an overestimation of the likely response to selection (McGlothlin and Galloway, 2014).

Maternal effects are often environment sensitive (Rossiter, 1998; Galloway, 2005; Räsänen and Kruuk, 2007; McAdam *et al*., 2014). It is unclear, however, whether challenging or favourable conditions will result in stronger maternal effects (Charmantier and Garant, 2005; Rowiński and Rogell, 2017). A harsher environment is typically assumed to increase variation among offspring from different mothers (Charmantier and Garant, 2005; Räsänen and Kruuk, 2007). This may, for example, stem from differences in maternal ability to provision offspring (Parichy and Kaplan, 1992). On the other hand, marine sticklebacks showed weaker maternal effects under stressful conditions (warmer temperature) (Shama *et al.*, 2014). Similarly higher population densities reduced offspring differences attributable to variation in maternal age in soil mites (Plaistow and Benton, 2009). Environment-dependent variation in maternal effects like this represents a form of Maternal-by-Environment interaction, M×E (Figure 1B). Another form of M×E occurs, irrespective of change in variance across environments, if individual mothers differ in how they affect their offspring in an environment-specific manner (Figure 1C). In an experiment with great tits for instance (Berthouly *et al.,* 2008), mothers supplemented with carotenoids raised smaller nestlings than the (un-supplemented) control mothers. When the brood size was increased, however, the result was opposite and the offspring of carotenoid-supplemented mothers grew faster compared to controls (Berthouly *et al*., 2008). A similar interaction was also observed for offspring immune response (Berthouly *et al*., 2008). The level of maternal carotenoids thus affects the fate and relative performance of the great tit offspring depending on the brood size. This interaction would cause cross-over of the maternal effects reaction norms because of the change in rank of maternal siblings between the two environments (Figure 1C). In general, a quantitative genetic analysis of potential M×E can help us to identify the mechanisms that increase similarity between siblings within the same environment beyond genetic relatedness. That analysis would be especially useful if such mechanisms themselves are undescribed, or their role is not fully understood. Many cases probably involve a mixture of changes in ranking and variances (as in Figure 1D), but the change in relative phenotype of offspring (i.e. shift in the rank) due to maternal effects is less often accounted for. In general, ignoring significant levels of M×E might bias estimates of additive genetic effects in a host-specific way and inflate G×E as shown by Vega-Trejo *et al.* (2018). This could partially explain inconsistency in G×E estimation across different studies (Saltz *et al.*, 2018).

**Figure 1.**
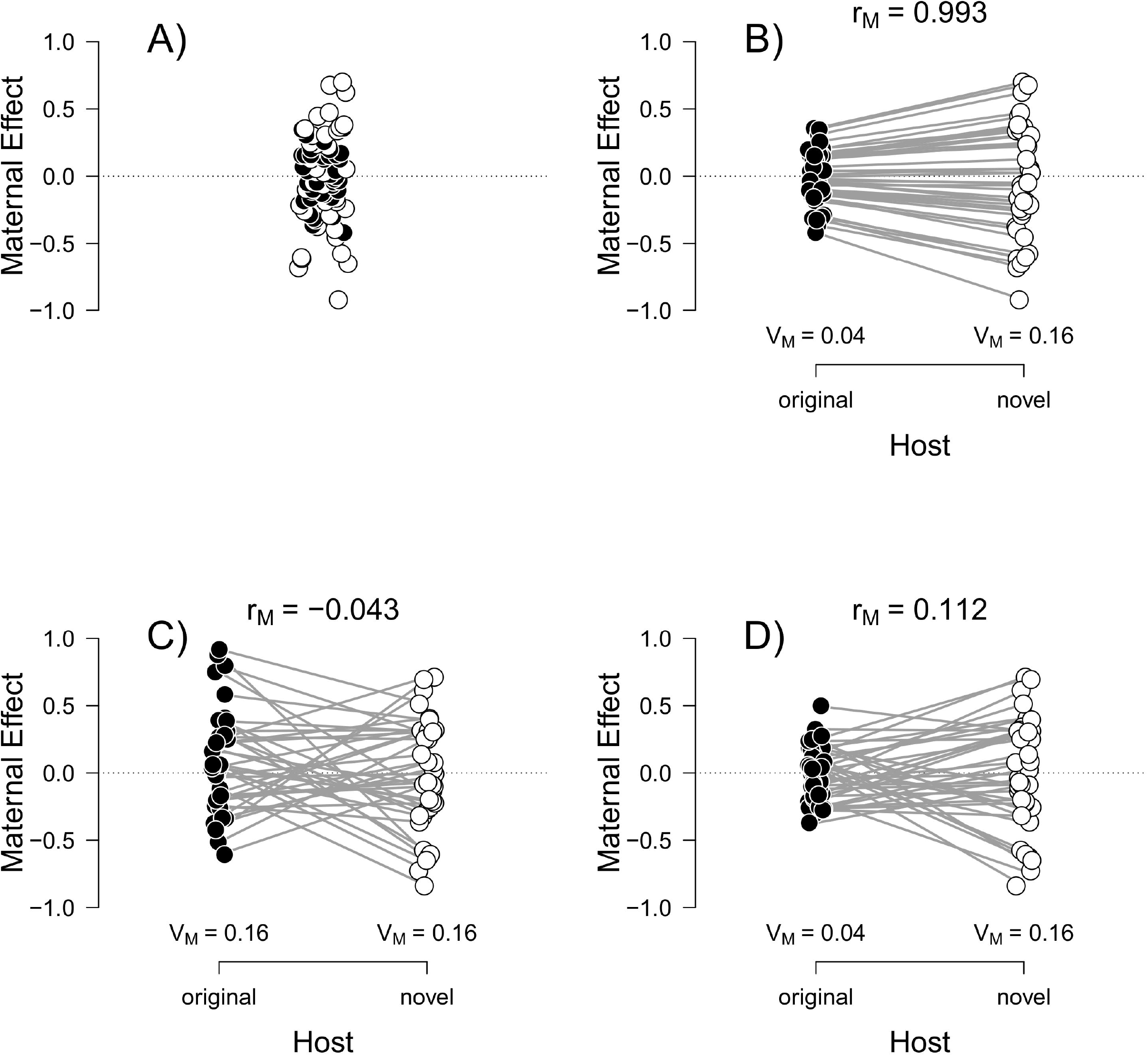
Illustration of how maternal effects might differ between environments (host beans in our study). Panel A) shows that ignoring the potential for environment-specific maternal effects, as indicated by the colour of points representing maternal effects in the original (black points) and the novel host (white points), could be misleading. Panel B) represents a scenario where variance due to maternal effects (V_M_) differs between the environments, i.e. the first type of Maternal-by-Environment interaction (M×E). Panel C) illustrates the other type of M×E where the rank of maternal effects differs between the hosts. The evidence for the shift in ranking comes from the low cross-environmental correlation in maternal effects (rM) despite the maternal effects variance (V_M_) being comparable between the hosts. Panel D) represents a combination of B) and C).

Phytophagous insects are a valuable model group to study life-history evolution, especially when interested in trade-offs arising from specialization to a preferred host (Via and Lande, 1985; Agrawal, 2000). At the same time, phytophagous insects exhibit a variety of maternal effects that influence key life-history traits (Fox and Dingle, 1994; Rossiter, 1996; Fox and Czesak, 2000; Van Asch *et al.*, 2010), including host-specific maternal effects (Fox *et al.,* 1997; Cahenzli and Erhardt, 2013). In the current study, we investigated the presence of maternal effects, and the potential for an interaction between maternal effects and offspring rearing host environment, on general offspring life-history traits, as well as daughters’ egg laying preferences for different host types in the seed beetle, *Callosobruchus maculatus* (Coleoptera: Chrysomelidae). This beetle is a common pest of stored legumes worldwide. The larvae feed on multiple legume species (family Fabaceae) with varying degrees of success (Gompert and Messina, 2016; Price *et al*., 2017; Messina *et al*., 2018). Females lay eggs on host beans shortly after copulation (Mitchell, 1975). Host choice is crucial as larvae cannot move between beans. Incorrect decisions about where a female chooses to lay her eggs inevitably lead to lower quality, or even dead, offspring (Mitchell, 1975; Messina and Fry, 2003; Messina *et al*., 2007). Larvae hatch 4-5 days after oviposition and burrow into the bean to feed on the endosperm. A single seed commonly harbours multiple larvae. The variable level of larval competition within seeds results in different sized adult beetles, which affects their condition, female reproductive performance and egg size (Fox and Savalli, 1998). Depending on the temperature and host species, adults typically emerge within 26-36 days of oviposition (Fox *et al*., 2003; Messina, 2004a). *Callosobruchus maculatus* shows large sexual dimorphism in life-history traits with males emerging earlier, being smaller and having a shorter lifespan than females (Guntrip *et al*., 1997; Fox, Bush, *et al*., 2004; Iglesias-Carrasco *et al*., 2020).

We used a full-sib/half-sib split brood design to tease apart the role of additive genetic and maternal effects on *C. maculatus* life-history traits when larvae develop on two host types – original (cowpea) and novel (mung bean). We hypothesised that:

1. There are strong maternal effects on offspring life-history traits (Fox, 1993; Messina and Fry, 2003). This led us to predict that a considerable component of offspring phenotypic variation, beyond that due to additive genetic effects, would be due to offspring having different mothers.
2. Maternal effects are environment-specific (M×E). Specifically, we predicted that the maternal effects variance would differ between offspring host environments (Figure 1B), and/or that the rank of offspring from the same mother will change between the two host types decreasing the cross-environmental maternal correlation (Figure 1C, D).
3. Novel host type will be more challenging for offspring. Hence, offspring developing in the novel host would suffer reduced performance compared to those developing in the original host type (i.e. lower larval survival, longer larval development, lower body mass at emergence, and shorter adult lifespan).
4. Maternal host preference will predict offspring performance. We predicted that offspring would perform better on the host type preferred by their mother when she laid her eggs.

## Methods

### Experiment overview

We aimed to estimate genetic and maternal effects for four life-history traits and host egg laying preference. To do this we used a half-sib/full-sib split brood design with offspring reared on two different host types – original or novel. We created the parental generation for our experiment by mating beetles from our stock population. We mated parents and tested mother egg laying preference for the two host types. We then let them lay eggs on both host types and measured offspring life-history traits and the daughters’ host preference. We used animal mixed-effects models to test our predictions.

### Origin and maintenance of stock beetles

Beetles used as the parental generation for our experiment were obtained from a large stock population originally sourced from the University of Western Australia (Perth, Australia) in 2017 where they had been bred on cowpea (*Vigna unguiculata,* Fabaceae) for at least 90 generations. We maintained this stock in our lab at the Australian National University on cowpea for another 9 generations before we began this experiment. Our stock was maintained in four, regularly mixed (every 5 generations) populations of over 500 individuals, each kept on cowpea at 25-26 °C.

### Establishing the parental generation (P)

To obtain virgin males and females for the parental generation (P), we exposed approximately 2000 un-infested cowpea beans to stock beetles for a period of 48 h. Each bean, which had 5-8 eggs on its surface (a density which is usual for our stock), was then placed in an individual Eppendorf tube (with a pinhole for airflow). Once isolated, we monitored these beans until adults began to emerge. We collected virgin beetles each morning and used them in our experiment on the same day. We knew that beetles were virgins as they were either the only beetle to have emerged that day, or all beetles that had emerged were of the same sex. Every evening, we discarded any extra beetles that had emerged during that day to aid in the collection of virgins the following day. Using the beetles that emerged each morning, we mated males and females according to a full-sib/half-sib breeding design: each male was sequentially mated with four random females over the day and the mating order was noted (similar to Fox *et al.* (2003)). Pairs that did not copulate within 30 min were separated for half an hour before being placed together for another mating attempt. Each female was weighed (to the nearest 0.001 mg) prior to mating (Cubis Ultra-Micro balance, Sartorius Lab Instruments GmbH., Goettingen, Germany). All matings took place over six days (‘day mated’, range 1-6, see *Model comparison and partitioning of phenotypic variance).* There is therefore a positive correlation between parental developmental duration (egg to adulthood) and ‘day mated’, but it is imperfect because parents emerged from eggs initially laid over two days (i.e. 48 hours; see above). There is no correlation between ‘day mated’ and adult parental age because all matings occurred on the day that the parents emerged.

### Maternal host preference

We used mung bean (*V. radiata*) as the novel host type. We define the host as novel in that it was ‘unlikely to have been experienced by the study population within an evolutionary timescale’ (Rowiński and Rogell, 2017), approximately 100 generations in our case (see also Kawecki (1995). Cowpea and mung are both suitable hosts for *C. maculatus* (Messina, 2004a; Fox and Messina, 2018), but populations kept on a specific host for multiple generations usually show better performance on their usual host than a novel host (Messina, 2004a).

Once females had mated, we conducted choice trials to determine their preferred host type for egg laying. We considered females ready to lay eggs once the pair dismounted (Wilson and Hill, 1989). For the host preference trials, we mixed cowpeas (original host) and mung beans (novel host) in covered Petri dish (ø 5.5 cm). We used proportionally more mung than cowpea beans (8:4) to roughly equalise the available surface area of each bean type (similar to 10:5 used by Messina and Slade (1997)). Cowpea beans have an approximately 1.6-times larger surface area (Paukku and Kotiaho, 2008) and are 4-times heavier than mung beans (in our study the average ± SD bean mass was 294±45 mg for cowpea, and 72±8 mg for mung bean). Females were left to lay eggs for two hours, after which they were removed and the number of eggs on each bean was counted. Relative preference was calculated as the proportion of the eggs that a female laid on the original host type (cowpea). The values for relative host preference therefore ranged between 0 and 1.

### Generation of offspring (F1)

Directly after the host preference trial, we transferred individual dams to plastic containers (ø 4 cm, height 6 cm) with 10-13 mung beans and left them to lay eggs for 14-18 h. We then moved dams to new containers with 10-13 cowpea beans for 9-10 h. This difference in laying time was required because initial trials showed that more time was necessary for females to lay a sufficient number of eggs on the novel host (mung beans), than on cowpea (the original host). Females were presented with mung beans first to prevent them from laying all their eggs on what we expected to be the preferred host type (cowpea) (Messina and Slade, 1997). Maternal age and/or laying order effects are unlikely to have influenced differences between mung and cowpea-reared offspring due to the short period of time we allowed females to lay eggs (i.e. <24 h). Previous studies have shown no effect of female age (hence laying order) on offspring quality during the first days of laying after emergence (Wasserman and Asami, 1985; Fox, 1993; Fox and Dingle, 1994; Iglesias-Carrasco *et al*., 2018). Further, one of the authors (MLH) found, in a separate experiment looking at oviposition preferences and offspring traits over the same time frame, that the order in which host beans are presented does not have any observable effect on offspring survival, development duration, or body mass at emergence (Zhang *et al*., in preparation).

Once females had laid eggs on both host types, we collected up to 10 beans of each host type for each female. If a female had laid eggs on fewer than 10 beans of a given type, we used them all. We ensured that each bean had only one egg laid on it by scraping off surplus eggs with a scalpel. We then weighed beans, to the nearest 0.001 mg, within 24 h of oviposition to measure the resources available to the larvae. Beans with an egg were then placed individually in Eppendorf tubes with perforated lids and incubated at 26 °C.

We started regular monitoring of F1 emergence on day 24 post-oviposition. However, we missed the onset of emergence of 40 beetles on mung (1.3% of all emerged offspring) by approximately 1 day as they emerged sooner than expected. Uncertainty in the date of their emergence could introduce noise into the estimates of larval development duration, body mass at emergence and adult lifespan, so we removed these beetles from the final dataset. Including these early emerging beetles did not qualitatively change our results.

### Measurement of offspring (F1) traits

We recorded the date of offspring emergence from the bean, sex and body mass at emergence (to the nearest 0.001 mg), and then removed the bean from the tube. After weighing, the beetle was returned to its tube and checked daily for survival. We also tested the host preference of two newly emerged female offspring per dam raised on each host type (total N = 4/dam). To do this we mated each daughter with a randomly selected male from the stock and then ran an egg laying preference trial identical to that described above for their mothers. We monitored survival until the death of all emerged offspring (day 79 post-oviposition). At that time, we also censused larval (egg-to-emergence) survival, assuming that larvae in beans from which a beetle had not yet emerged had died.

### Sample sizes

Our target sample size was determined *a priori* based on Lynch and Walsh (1998). We created 89 families (89 sires with 356 dams) and aimed for 10 offspring per dam, per host type (i.e. 7 120 offspring). The final sample size (N = 3 431) is lower due to unsuccessful matings or too few eggs being laid. Despite not reaching our target sample size our final sample size is still higher than that in other similar studies of maternal effects (the studies in a meta-analysis by Moore *et al.* (2019), for example, have a mean sample size of 1 370, and a much lower median of 555). We analysed data for sons and daughters separately due to their large sexual dimorphism which invariably leads to strong interactions between sex and other effects. We excluded offspring of dams that did not lay eggs during host preference trials (285 individuals) and data from 13-24 daughters and 12-27 sons with missing, or extreme values (>3 standard deviations (SD) from the mean), with the exact number depending on the focal trait. We analysed four offspring life-history traits: (egg-to-emergence) ‘larval survival’ (dead or emerged; N = 3 146), ‘duration of larval development’ (number of days between oviposition and offspring emergence; N = 1 420 daughters and 1 384 sons), ‘body mass’ (weight at emergence in mg; N = 1 431 daughters and 1 399 sons) and ‘adult lifespan’ (number of days between offspring emergence and death; N = 1 425 daughters and 1 389 sons). In addition, we measured the host egg laying preferences of 665 daughters. The analysed offspring came from 82 sires and 245-249 dams for the life-history traits, and 239 dams for host preference analysis. The proportion of offspring reared on the original host type was 52-57 % (see Supplementary Table S1). Prior to any statistical analysis, but after all data had been collected, we registered this project on the Open Science Forum webpage: https://osf.io/ft7eq/?view_only=0bab0a33bb4246adb64c919601a72757 The final analytical approach we used was different, however, from the original plan – partly due to feedback from reviewers. We explain our changes in the *Annotated registration* section in the Supplementary Information.

### Overview of data analysis approach

Our main objective was to test for maternal effects on the offspring traits (Prediction 1) and their potential interaction with offspring rearing environment (Prediction 2). First, we built a ‘minimal model’ containing only fixed effects directly attributable to the experimental design. To test whether maternal effects are important, we then created a set of candidate models with different random effects structures (Table 1). We used the best-selected model for partitioning of phenotypic variance (Table 2). Finally, we assess the effect of host type (Prediction 3) and dam host egg laying preference (Prediction 4) on offspring traits with a ‘full’ model containing additional fixed effects (Table 3).

**Table 1.** Model comparison to estimate the best random effects structure for minimal animal MCMC models. The table shows the difference between Deviance Information Criterion (DIC) for each candidate model and the model with the lowest DIC (denoted as ‘-’). Model ‘G’ contained additive genetic variance only. The model with ‘M’ included also maternal effects variance. Models where variance was estimated separately ‘sep’ for each host type are ‘G_sep_’ or ‘M_sep_’. Models ‘G_cov_’ (or ‘M_cov_’) estimated covariance ‘_cov_’ between the two host types in addition to the hostspecific variance. When the host-specific interaction was fitted as a random effect (models ‘_sep_’ and ‘_cov_’), we also estimate residual variance separately for each host type. Further details on model fitting are in the Methods. Please note that the low DIC for model ‘G_cov_’ means that the genetic covariance between the host types is different from zero. It is, however, inconclusive about Genotype-by-Environment interaction as the covariance can still be highly positive (i.e. the rank order of genotypes does not change). So we also examined the cross-environmental genetic correlation (r_G_) estimates from the model given in Table 2. DIC is not reliable for model comparison in non-Gaussian traits (Hadfield, 2010; Wilson *et al*., 2010) and we thus did not perform model selection for larval survival and host egg laying preference of daughters and used model G_cov_ for inference.

| Model | Random effects | Development duration |  | Body mass |  | Adult lifespan |  |
| --- | --- | --- | --- | --- | --- | --- | --- |
|  |  | daughters | sons | daughters | sons | daughters | sons |
| G | animal | 137.6 | 225.4 | 82 | 78.8 | 70.4 | 125.6 |
| G+M | animal + dam | 151.8 | 233.3 | 136.3 | 115.9 | 79.4 | 153.2 |
| G<sub>sep</sub> | idh(HOST):animal | 10.4 | - | 18.1 | 46.3 | 15.6 | 72.8 |
| G<sub>cov</sub> | us(HOST):animal | - | 0.4 | - | - | - | - |
| G<sub>sep</sub>+M<sub>sep</sub> | idh(HOST):animal + idh(HOST):dam | 42.5 | 29.6 | 97.5 | 108.9 | 39.2 | 133.1 |
| G<sub>cov</sub>+M<sub>sep</sub> | us(HOST):animal + idh(HOST):dam | 23.7 | 33.5 | 31.3 | 22.1 | 16.3 | 17.6 |
| G<sub>cov</sub>+M<sub>cov</sub> | us(HOST):animal + us(HOST):dam | 30.1 | 31.3 | 53.3 | 63.6 | 17.2 | 34.6 |

**Table 2.** Phenotypic variance partitioning based on the best model from the model comparison. We used model ‘G_cov_’ for all the traits (Table 1) as it estimates cross-environmental genetic covariance and models with DIC difference <2 are usually considered equally supported. We estimated host-specific additive genetic effects (V_A_), proportion of additive genetic variance in total phenotypic variance (V_P_) – heritability (h^2^), additive genetic variance standardized over trait mean – evolvability (CV_A_), and cross-environmental genetic correlation (r_G_). The estimates and their credible intervals (in brackets) come from posterior distribution of sex-specific animal MCMC models.

| Host type | Par. | Development duration |  | Body mass |  | Adult lifespan |  |
| --- | --- | --- | --- | --- | --- | --- | --- |
|  |  | daughters<br>(N = 1 420) | sons<br>(N = 1 384) | daughters<br>(N = 1 431) | sons<br>(N = 1 399) | daughters<br>(N = 1 425) | sons<br>(N = 1 389) |
| original | $V_A$ | 0.430 (0.105,0.678) | 0.448 (0.208,0.876) | 0.215 (0.147,0.317) | 0.070 (0.034,0.114) | 19.981 (11.423,26.851) | 9.694 (5.578,14.841) |
| | $V_P$ | 1.768 (1.578,1.948) | 2.167 (1.979,2.409) | 0.501 (0.446,0.555) | 0.310 (0.282,0.343) | 39.267 (35.595,45.099) | 28.298 (24.908,31.138) |
| | $h^2$ | 0.229 (0.084,0.387) | 0.210 (0.082,0.374) | 0.447 (0.327,0.615) | 0.225 (0.122,0.358) | 0.459 (0.301,0.624) | 0.333 (0.218,0.505) |
| | $CV_A$ | 0.020 (0.012,0.027) | 0.021 (0.014,0.030) | 0.084 (0.066,0.097) | 0.075 (0.058,0.100) | 0.195 (0.148,0.226) | 0.184 (0.146,0.234) |
| novel | $V_A$ | 0.349 (0.213,0.622) | 0.527 (0.355,0.892) | 0.250 (0.176,0.319) | 0.103 (0.068,0.148) | 16.664 (10.973,27.715) | 16.304 (11.261,24.817) |
| | $V_P$ | 0.956 (0.853,1.083) | 1.154 (1.028,1.339) | 0.374 (0.340,0.426) | 0.211 (0.186,0.236) | 37.645 (33.595,43.241) | 26.675 (23.763,31.265) |
| | $h^2$ | 0.365 (0.241,0.606) | 0.495 (0.348,0.708) | 0.649 (0.501,0.786) | 0.513 (0.355,0.669) | 0.507 (0.334,0.684) | 0.627 (0.480,0.854) |
| | $CV_A$ | 0.020 (0.016,0.026) | 0.026 (0.020,0.031) | 0.077 (0.065,0.087) | 0.082 (0.067,0.099) | 0.168 (0.131,0.204) | 0.208 (0.176,0.260) |
| | $r_G$ | 0.706 (0.277,0.981) | 0.147 (-0.279,0.533) | 0.901 (0.670,0.995) | 0.996 (0.750,1.000) | 0.790 (0.615,0.982) | 0.967 (0.777,0.999) |

**Table 3.** The effect of host type, other experimental variables and dam host preference on three offspring life-history traits (fixed-effects estimates from the full animal models). The intercept always corresponds to the trait average on the original host type and the ‘novel HOST effect’ is the effect of the novel host type (i.e. difference between the two means). The same applies to interaction terms, where the interaction gives the estimate for the difference between the effect on the novel and the original host type. We embolden any credible intervals (in brackets) that do not overlap zero. The estimates and their credible intervals come from posterior distribution of sex-specific animal MCMC models.

| Term | Development duration [days] |  | Body mass [mg] |  | Adult lifespan [days] |  |
| --- | --- | --- | --- | --- | --- | --- |
|  | daughters<br>(N = 1 420) | sons<br>(N = 1 384) | daughters<br>(N = 1 431) | sons<br>(N = 1 399) | daughters<br>(N = 1 425) | sons<br>(N = 1 389) |
| Intercept (original HOST) | 31.899 (31.713,32.103) | 31.442 (31.187,31.607) | 5.784 (5.660,5.904) | 3.466 (3.414,3.593) | 23.62 (22.432,24.739) | 16.298 (15.338,17.259) |
| novel HOST effect | **-1.285 (-1.388,-1.115)** | **-1.148 (-1.296,-0.935)** | **0.728 (0.650,0.792)** | **0.349 (0.303,0.408)** | **3.338 (2.595,3.975)** | **2.355 (1.792,2.885)** |
| bean mass (original) | **0.145 (0.035,0.229)** | 0.105 (-0.017,0.189) | **-0.083 (-0.130,-0.032)** | **-0.039 (-0.089,-0.011)** | **-0.499 (-0.930,-0.111)** | **-0.365 (-0.742,-0.038)** |
| dam mating order | 0.060 (-0.002,0.136) | 0.054 (-0.013,0.134) | 0.007 (-0.037,0.052) | 0.006 (-0.017,0.047) | -0.306 (-0.766,0.069) | 0.210 (-0.116,0.587) |
| day mated (original) | **0.326 (0.206,0.454)** | **0.122 (0.028,0.291)** | 0.058 (-0.006,0.149) | **0.074 (0.037,0.139)** | 0.443 (-0.113,1.273) | **0.797 (0.199,1.262)** |
| dam preference for<br>cowpea (original) | -0.073 (-0.180,0.031) | -0.025 (-0.174,0.068) | **0.086 (0.027,0.144)** | 0.000 (-0.042,0.046) | 0.219 (-0.366,0.733) | 0.128 (-0.267,0.572) |
| Interaction: bean mass<br>(novel) | -0.092 (-0.196,0.032) | 0.031 (-0.131,0.138) | 0.047 (-0.022,0.102) | 0.040 (-0.011,0.091) | 0.456 (-0.052,1.124) | -0.060 (-0.463,0.560) |
| Interaction: day mated<br>(novel) | -0.017 (-0.141,0.142) | 0.112 (-0.026,0.326) | -0.057 (-0.127,0.020) | -0.018 (-0.072,0.035) | -0.625 (-1.272,0.117) | -0.360 (-0.910,0.237) |
| Interaction: dam<br>preference (novel) | 0.051 (-0.056,0.206) | 0.051 (-0.084,0.228) | **-0.081 (-0.146,-0.015)** | 0.008 (-0.038,0.063) | -0.369 (-0.916,0.402) | -0.109 (-0.590,0.470) |

### Model specification, model comparison and partitioning of phenotypic variance

We started data analysis with a minimal mixed-effects model that contained, in the fixed-effects part, only variables ensuing from the experimental design. These were ‘host type’ (original/novel), ‘bean mass’ (standardized to zero mean and 1 SD variance, within each host type, to look at the effect of size within each host type), ‘dam mating order’ (i.e. dam position in a sire’s mating sequence, integer 1 to 4), and ‘day mated’ (integer 1-6). We did not fit ‘day mated’ as a random effect because of the low number of levels (Bolker *et al*., 2009). We specified the random effects structure using an animal model approach (Kruuk, 2004; Wilson *et al*., 2010), where variance-covariance estimates between relatives are computed from a pedigree.

We fitted the models using Bayesian framework with package MCMCglmm (ver. 2.29) (Hadfield, 2010) in R software (ver. 4.0.0; R Core Team, 2020). The continuous response variables (duration of larval development, body mass and adult lifespan) were fitted using Gaussian family and identity link, while larval survival (dead or emerged) was fitted using ‘categorical’ family (binomial distribution with logit link). Daughter host preference was treated as a bivariate vector with the number of eggs on the original host and the total number of eggs collected in the egg laying preference trial using ‘multinomial2’ family (binomial distribution and logodds ratio link function) (Hadfield, 2010).

To test for the best random effects structure, we performed model comparison on a set of 7 candidate models (Table 1) for each trait using the Deviance Information Criterion (DIC) (Spiegelhalter *et al*., 2002). DIC is a Bayesian alternative to Akaike Information Criterion (AIC) that takes into account the fit of the model but penalizes for model complexity, with lower DIC values indicating a better model (Hadfield, 2010; Wilson *et al*., 2010).

Model comparison with DIC should be performed cautiously for nonGaussian response variables (Hadfield, 2010; Wilson *et al.*, 2010), so we did not conduct model selection for larval survival or host preference.

The simplest model ‘G’ included only additive genetic effects specified by the random effect of ‘animal’. We then also included dam identity to estimate maternal effects in addition to additive genetic effects (‘G+M’ model). These two models do not, however, take into account possible hostspecific differences in respective variance components (this is shown in Figure 1A). We thus formulated more realistic models that estimated the variance component (including residual variance) separately for each host type (as in Figure 1B). These were specified through random-effect interactions (Hadfield, 2010) with host type ‘idh(HOST):animal’ for ‘G_sep_’, or with ‘idh(HOST):dam’ for ‘M_sep_’ models (Table 1). This is similar to fitting two separate host-specific models. To account for potential change in the ranking of additive genetic or maternal effects (Figure 1C), we fitted models that also estimated cross-environmental covariance (‘G_cov_’ or ‘M_cov_’). When specifying variance-covariance matrix of the random-effect interaction we used ‘unstructured’ form, e.g. ‘us(HOST):animal’ (Hadfield, 2010). As a result, we not only obtained separate variance estimates for each host type, but also the corresponding covariance to compute the cross-environmental correlations (Figure 1D) – the proxies for Genotype-by-Environment (G×E), or Maternal-by-Environment (M×E) interactions (Lynch and Walsh, 1998). In the models estimating separate variances per host type, we always specified host-specific residual variance (as ‘rcov=~idh(HOST):units’) (Hadfield, 2010).

We then performed phenotypic variance partitioning based on the outputs of the best-selected minimal model for each trait. We calculated the maternal-effects proportion (m^2^), narrow-sense heritability (h^2^), evolvability (coefficient of additive genetic variation, CV_A_) and the cross-environmental correlation for additive genetic (r_G_ – potential G×E interaction) and for maternal effects (r_M_ – potential M×E interaction). We used maternal effects proportion, 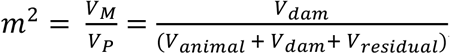, to quantify the importance of maternal effects (as in e.g. Messina and Fry (2003); Vega-Trejo *et al.* (2018); Moore *et al.* (2019)). The V_M_ contains both genetic and non-genetic maternal effects, and potentially also genetic dominance, as we did not separate them with our experimental design. We calculated heritability as 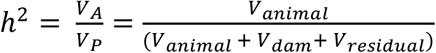 (Lynch and Walsh, 1998), assuming epistatic effects were negligible. Evolvability was defined by (Houle, 1992) as 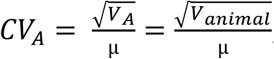, where μ is the mean of a trait. Evolvability standardizes additive genetic variance over the trait mean and is therefore useful to compare the potential evolutionary response among traits, unlike heritability which is conditional on the amount of residual phenotypic variation (Hansen *et al*., 2011). The correlation between additive genetic effects on the two host types (G×E interaction) was 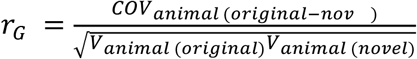. The same formula was used for maternal effects correlation between the two hosts (potential M×E interaction) using the dam-related covariance across the two host types. Estimates of V_A_, V_P_ and h^2^ for larval survival and host preference were back-transformed to the original data scale using the function ‘QGparams’ (from QGglmm package, ver.0.7.4; de Villemereuil *et al.* (2016)).

### Testing for host effect and dam host egg laying preference on offspring phenotype

To determine the effect of novel host type on offspring performance (Prediction 3) and to test whether offspring performed better on the host type that was preferred by their mother (Prediction 4), we built a ‘full model’. The full mixed-effects model always included all the terms outlined above for the minimal model with the random effects from the best-selected minimal model (Table 1). We then included ‘dam host preference’ (i.e. the relative preference ratio: 0-1 values) and the interaction between preference and host type ‘HOST:dam host preference’ to test if the effect of the strength of the preference for the original host on offspring traits differs depending on the host type. We also included interactions between host type and bean mass ‘HOST:bean mass’, as well as host type and day mated ‘HOST:day mated’ as fixed effects to test for potential hostspecific effects.

### Model fitting

To fit the MCMC models, we used non-informative priors for the random-effects variance (G) with expected variance-covariance matrix at limit ‘V’ = 1, and the degree of freedom ‘nu’ = 0.002 for each random term (with nu = 1.002 in models with the random effects estimated separately per host type) (Wilson *et al*., 2010). To further improve mixing and effective sample size, we specified parameter-expanded priors as ‘α mu’ = 0 and ‘α V’ = 1 000 (Hadfield, 2010). Priors for residual variance (R) were set as V = 1 and nu = 0.002 (nu = 1.002 when residual variance was estimated separately for each host type). In the analysis of survival, the residual variance had to be fixed to 1, so that the binomial mixed-effects model could be estimated (Hadfield, 2010). For fixed effects (B), we retained default priors with mean = 0 and variance = 10^10^. To generate posterior distributions from the minimal and full models, we ran 360 000 iterations with burn-in of 60 000 and thinning interval of 300, so that we obtained an effective sample size of 1 000 for each estimated model term. For larval survival, we increased the iterations to 5 500 000 with a burn-in of 500 000 and thinning interval of 5000 as we were aiming for an autocorrelation <0.1 at the lag corresponding to the thinning interval (Wilson *et al*., 2010). Model convergence was evaluated by visual examination of the traces using function ‘plot(mcmc.model)’. We assessed the importance of individual model terms based on their credible intervals (CrI).

## Results

### Prediction 1 and 2 – testing for maternal effects and M×E interaction

The model comparison suggests very weak maternal effects for all the measured traits. The models incorporating maternal effects performed poorly (Table 1), and the best model contained only additive genetic effects estimated for each host type and the cross-environmental covariance (the ‘Gcov’ model). This shows that either additive genetic variance, residual variance or both differed between host types and that the cross-environmental genetic covariance differed from zero. Estimate of the cross-environmental covariance has to be examined directly to see whether the correlation differs from +1 indicating a potential Genotype-by-Environment interaction (see below).

### Components of phenotypic variance related to additive genetic effects – heritability, evolvability and G×E

We observed higher heritability for offspring life-history traits in the novel host type (h^2^ = 0.365-0.649), than on the original host (h^2^ = 0.210-0.459) (Table 2). However, the additive genetic variance was substantially higher in the novel host only for adult lifespan of sons. For the other traits, we recorded a similar magnitude of additive genetic variance in the two hosts, but higher residual phenotypic variance in the original host type, thereby reducing trait heritability (Table 2). The presence of similar levels of additive genetic variance in both hosts is also apparent in the estimates of evolvability (variance due to additive genetic effects standardised over trait means) which were comparable for the two host types (Table 2). When comparing evolvability among traits (across sexes and host types) we found that it was highest for adult lifespan (CV_A_ = 16.8-20.8 %, Table 2). For the non-Gaussian traits (larval survival and host egg laying preference), estimates of additive genetic variance and heritability obtained from the Gcov model were very low (h^2^ = 0.001-0.060, Table 4A) when compared to the Gaussian life-history traits.

**Table 4.** Phenotypic variance partitioning (A) and full model inference (B) based on G_cov_ model for the non-Gaussian traits – larval survival and host egg laying preference. The estimates for the variance parameters (A) are back-transformed to original data scale. The effect of host type, the additional experimental variables and dam host preference (B) for larval survival is on logit scale and for host preference on log scale. We embolden any credible intervals (in brackets) that do not overlap zero. The intercept always corresponds to the trait average on the original host type and the ‘novel HOST effect’ is the effect of the novel host type (i.e. difference between the two means). The same applies to interaction terms, where the interaction gives the estimate for the difference between the effect on the novel and the original host type. The estimates and their credible intervals come from posterior distribution of animal MCMC models.

| Host type | Par. | Larval survival | Host preference |
| --- | --- | --- | --- |
|  |  | juveniles<br>(N = 3 146) | daughters<br>(N = 665) |
| original | $V_A$ | 0.006 | <0.001 |
| | $V_P$ | 0.094 | 0.671 |
| | $h^2$ | 0.060 | 0.001 |
| | $CV_A$ | 0.084 | 0.033 |
| novel | $V_A$ | 0.003 | 0.001 |
| | $V_P$ | 0.065 | 0.522 |
| | $h^2$ | 0.040 | 0.003 |
| | $CV_A$ | 0.055 | 0.071 |
| $r_G$ | | 0.286* (-0.469,0.985)* | -0.046* (-0.905,0.948)* |

\* these estimates are on the transformed (latent) scale

B)
| Term | Larval survival | Host preference |
| --- | --- | --- |
|  | juveniles | daughters |
|  | (N = 3 146) | (N = 665) |
| Intercept (original HOST) | 2.868 (2.341,3.441) | -0.409 (-0.528,-0.338) |
| novel HOST effect | 0.512 (-0.094,1.447) | **-0.247 (-0.324,-0.180)** |
| bean mass (original) | -0.161 (-0.353,0.050) | 0.010 (-0.046,0.049) |
| dam mating order | -0.021 (-0.206,0.124) | -0.001 (-0.027,0.047) |
| day mated (original) | 0.095 (-0.153,0.350) | -0.053 (-0.094,0.004) |
| dam preference for cowpea (original) | -0.042 (-0.258,0.183) | 0.014 (-0.041,0.057) |
| Interaction: bean mass (novel) | -0.035 (-0.376,0.221) | 0.017 (-0.075,0.076) |
| Interaction: day mated (novel) | -0.122 (-0.501,0.240) | -0.016 (-0.066,0.083) |
| Interaction: dam preference (novel) | 0.242 (-0.106,0.549) | -0.037 (-0.096,0.049) |

We recorded highly positive correlations between offspring performance on the two host types for additive genetic effects (rG) (Table 2). The only exception was for the development duration of sons, which exhibited only a weak genetic correlation between hosts (Table 2). The credible interval (−0.279, 0.533) suggests a Genotype-by-Environment interaction (G×E).

### Predictions 3 and 4 – testing for host type suitability and host preferenceperformance

Unexpectedly, given our predictions, offspring generally performed better on the novel host (mung) than on the original host (cowpea) (Table 3; Supplementary Table S1). We recorded very high larval survival overall, however, so that survival on mung beans was only marginally better than on cowpea (Table 4B). We did not find a positive relationship between the strength of the dam host preference for egg laying and the performance of their offspring on either host type. The one exception was for daughter body mass: daughters of dams that more strongly preferred the cowpea emerged heavier when raised on cowpea, but not when raised on mung beans (Table 3). Daughters reared on the original cowpea host also exhibited a higher egg laying preference for cowpeas than those reared on mung beans. Raw average host preference ratio of daughters reared on cowpea was 0.681 versus 0.517 for those from mung (where 1 is a strict preference for cowpeas). The host preference ratio of their mothers (reared on cowpea) was 0.724.

### Effect of experimental variables on offspring traits – bean size, mating date and mating order

Host-specific relative bean size (zero-centered per host type) and mating date were important predictors of offspring traits. For both host types, offspring that developed in larger beans had a poorer outcome: increased development duration in daughters, as well as reduced body mass and lower adult lifespan in both sexes. The effects of mating day on offspring performance were, however, mixed (Table 3). Offspring of later mated families took longer to develop, but sons emerged at a larger size and lived longer. Daughters from later mated families also showed a modest decrease in the preference for the original host type (Table 4B). Finally, if a dam was later in the sire’s mating sequence this marginally extended the development duration of her daughters.

## Discussion

### Summary

Our main aim was to test for environment-specific maternal effects on offspring life-history traits and their correlation, that is, Maternal-by-Environment interactions (M×E). We used a full-sib/half-sib split brood design with seed beetles (*C. maculatus)* reared on two types of host: original – cowpea; and novel – mung bean. Maternal effects, however, proved to be negligible for all of the measured traits, on both host types. Instead, we found that additive genetic variance played an important role in determining phenotypic variation. Contrary to our predictions, offspring reared on the novel host type performed better for all the measured traits. Daughters whose mothers preferred to lay eggs on the original host were heavier on this host than those of mothers that preferred the novel host type. There was, however, no equivalent effect of host preference on body mass in the novel host. Below we discuss our findings in more detail.

### Weak maternal effects

We used an original and a novel host type to examine whether mothers affect their offspring’s phenotype in a hostspecific way. Previous studies have shown that environmental stress can either increase or decrease the magnitude of maternal effects (Charmantier and Garant, 2005; Rowiński and Rogell, 2017). Maternal effects variation is sometimes not evident in resource rich conditions as differences among mothers are reduced compared to a poor environment where differences are exacerbated (e.g. Parichy and Kaplan (1992)). If both host types in our study are resource rich (especially when few eggs are laid per bean) this could explain the negligible maternal effects that we see here. Previous quantitative genetic studies employing a similar approach to ours have reported significant maternal effects in *C. maculatus.* This has been, however, in other hosts (adzuki *(Vigna angularis)* in Fox (1993)) or when testing the effect of bean removal after beetle emergence on adult traits (Messina and Fry, 2003). The host species we used are both thought to be highly favourable to *C. maculatus* (Messina, 2004a; Fox and Messina, 2018) (discussed further below). This, in addition to the fact we prevented larval competition with our experimental design, may have substantially reduced the role of maternal effects in the current experiment.

### Heritability was higher in the novel host type, but evolutionary potential was similar in both hosts

Heritability of fitness-related traits is traditionally assumed to be low, as additive genetic variation is expected to be depleted by natural selection (Mousseau and Roff, 1987; Hill, 2010). Egg size, body size, fecundity or lifespan were nevertheless previously shown to have intermediate to high heritability in *C. maculatus* (h^2^ from 0.27 to 0.74) (Fox, 1993; Messina, 1993; Fox *et al*., 2003; Fox and Messina, 2018). Here, we recorded considerable heritability in all life-history traits (except larval survival), with the highest values occurring for daughter body mass and son adult lifespan (both h^2^ > 0.65). Heritability was notably higher in the novel host environment (mung bean), with the one exception of adult lifespan of daughters. Holloway *et al.* (1990) proposed that a novel environment should yield higher additive genetic variance due to new genes being expressed thereby uncovering cryptic genetic variation (Hoffmann and Merilä, 1999). Indeed, Messina and Fry (2003) found that *C. maculatus* exhibited increased additive genetic variance for longevity under a novel environment (absence of host beans after emergence). Our results indicate, however, that it was lower residual variance in the novel host (mung bean), rather than increased additive genetic variation, that was responsible for higher heritability in the novel environment. Residual phenotypic variation may entail non-additive genetic, environmental variance and developmental noise (Rowiński and Rogell, 2017) and the influence of at least some of these factors apparently declined in the novel host type. More elaborated breeding design would be required (see e.g. Tucić and Šešlija (2007), or Bilde *et al.* (2008)), however, to examine their respective effects in *C. maculatus.* Interpretation of heritability as the proportion of additive genetic versus total variance can therefore become problematic (Hansen *et al*., 2011) and evolvability offers a more conclusive comparison of evolutionary potential (Houle, 1992; Hansen *et al.*, 2011). Similar values of trait evolvability indicated that the traits measured in our study possess similar evolutionary potential in the two host types.

Genetic trade-offs are thought to prevent populations of parasites or herbivores from becoming ‘masters of all trades’ and performing well on multiple hosts (Joshi and Thompson, 1995; Agrawal *et al*., 2010). In phytophagous insects, performance trade-offs on different hosts are, however, rarely reported (Ueno *et al*., 2003; Scheirs *et al*., 2005; Messina and Durham, 2015). This includes studies of *C. maculatus* where offspring from the same family usually perform similarly across hosts without exhibiting a G×E interaction (Fox, 1993; Guntrip and Sibly, 1998; Messina, 2004a). We also found strong positive cross-environmental genetic correlations indicating parallel reaction norms for offspring from different families when reared on the two host types. Only the development duration of sons showed a low genetic correlation between the two host types indicating that it could evolve differently on cowpea and mung (Via and Lande, 1985). It is, however, important to bear in mind that breeding design studies cannot definitively conclude there are no genetic trade-offs in adaptation to different hosts because they lack mechanistic explanation. This can be obtained through analysis of long-term selected lines (Agrawal *et al.*, 2010) or data on the potential genes involved (Gompert and Messina, 2016).

### Offspring performed better on the novel host type

We treated mung bean as a novel host type. This was despite mung bean regularly being used to rear other strains of *C. maculatus* (e.g. the Southern India strain (Fox, Bush, *et al*., 2004)), and regarded with cowpea as a favourable host species for *C. maculatus* (Fox and Messina, 2018). Our stock population was kept on cowpea for at least 100 generations. In addition, cowpea strains are overall more viable than those from mung bean (Messina, 2004a) and the cowpea is generally considered to be the ancestral host for *C. maculatus* (Kébé *et al*., 2017). Due to these reasons and the fact that a previous study showed that long-term maintenance on a specific host yields decreased viability on other hosts (Messina, 2004a), we predicted a lower suitability of the novel host type (mung bean). Our finding that offspring performed better on the novel host type (mung bean) was therefore unexpected. While offspring survival was almost 90 % on cowpea, it was even higher on the novel host (93 % on mung bean). There was no obvious cost to the offspring of rearing in the novel host, as the duration of larval development was shorter on mung bean. However, beetles that emerged from mung beans were larger and lived longer as adults than those that emerged from cowpeas. It is not straightforward, however, to tell which of the two host species is generally better for *C. maculatus,* because it depends on other factors such as the level of competition (Messina, 2004a; Paukku and Kotiaho, 2008; Fox and Messina, 2018), temperature (Stillwell *et al.*, 2007) and the population’s evolutionary history (Messina, 2004a). In the present study, we eliminated effects of larval competition by rearing one larva per bean. We therefore removed the fitness advantage that cowpea may provide due to its larger size when there is larval competition (Fox and Savalli, 1998). Surprisingly, relative bean size within a given host type also negatively affected body mass and lifespan as well as extending the developmental duration of daughters.

We cannot reject the hypothesis that the host effect recorded in our experiment was confounded by egg laying order, as eggs were laid first on mung bean (see Methods for the experimental justification). The lower performance of offspring on cowpea might have been because, following the preference trial, females were first provided with mung beans to lay eggs, and only after that with cowpeas. There are, however, several reasons to think that an effect of laying order in our experiment is unlikely. The lifetime fecundity of *C. maculatus* is 60-80 eggs (Mitchell, 1975; Credland and Wright, 1989; Wilson and Hill, 1989; Messina and Fry, 2003). The dams in our experiment laid around 30 eggs per 24 h and probably still possessed more than half of their lifetime fecundity after they had finishing laying in our experiment. Crucially, past studies suggest there is no negative effect of maternal age on offspring performance on the first day of egg laying (i.e. the relevant time period in our study) (Wasserman and Asami, 1985; Fox, 1993; Fox and Dingle, 1994; Iglesias-Carrasco *et al*., 2018; Zhang *et al*., in preparation). We therefore think that the sequence of host types available for egg laying is unlikely to explain the poorer offspring performance on cowpea.

### Host preference of *C. maculatus* increased daughter body mass and rearing host affected host preference

An egg laying preference for a particular host often predicts offspring performance on that host in phytophagous insects (Gripenberg *et al*., 2010). However, despite the critical role of oviposition choice for larval success in *C. maculatus,* of the five offspring traits we measured, the strength of dam preference for cowpea only correlated with daughter body mass. Dams that showed a stronger preference for cowpea produced heavier-than-average daughters on cowpea, but not when laying on mung. This contributes to mother fitness through higher daughter reproductive potential in cowpea as body size is a good proximate measure for fecundity in *C. maculatus* (Messina and Fry, 2003). In mung bean, however, daughters’ body mass was not affected by their mothers’ host preference and the preference-performance relationship thus only holds for daughter body mass in cowpea. The weak overall relationship between mother’s host preference for one host type and the performance of her offspring on that host suggests that *C. maculatus* females might aim to maximize their own fitness and not that of individual offspring (i.e. they prioritise offspring number over offspring quality), perhaps through saving searching time for the best host (Wood *et al.*, 2018).

Previous studies show that strains of *C. maculatus* maintained for > 40 generations on mung bean or cowpea exhibit a strong preference for their recent host type (Messina, 2004b). Fox, Stillwell, *et al.* (2004) used different host strains to demonstrate that switching a strain to another host for just a single generation has no effect on host preference. We found that while daughters reared on cowpea preferred cowpea over the novel host (mung bean), those reared on mung beans showed no preference for either host type. This suggests that there might be a G×E interaction affecting host egg laying preferences, but tracking of more generations is necessary to test whether the host preference evolves after the host switch (as in Fox, Stillwell, *et al.* (2004), or Messina (2004b)).

### Conclusion

Maternal effects were low for all measured traits in our study for offspring reared on either a novel or a familiar host. A next step in the exploration of whether M×E interactions affect estimates of genetic variation, and of G×E interactions specifically, would be to conduct experiments using less favourable novel hosts of *C. maculatus*, for example adzuki bean (Fox, 1993) or lentil (Gompert and Messina, 2016). The host species we used are both considered to be of high quality (Fox and Messina, 2018). By creating more challenging conditions for offspring, maternal effects might emerge as an important host-specific factor that affects variation in offspring performance (Fox *et al*., 1997; Räsänen and Kruuk, 2007). Despite the fact that maternal effects were negligible in our study, we still argue that it is important testing for M×E interactions. The M×E effects have potential to bias measurements of G×E and the lack of research on M×E interaction is an obstacle towards understanding their role in adaptive processes.

## Acknowledgements

We appreciate detailed comments by three anonymous referees that helped us to improve the manuscript. We thank Timothée Bonnet and Loeske E. B. Kruuk (RSB ANU) for very helpful discussions about phenotypic variation partitioning and Bayesian mixed-effect models. We are thankful to Carolina Dolan, Krish S. Sanghvi and Elroy Kwan Au (RSB ANU) for their help with the experiment. MV and PJCC were both supported by 6-month Endeavour fellowships from the Australian Government.

## Author contribution

MV, PJCC, MLH and MDJ conceived and designed the study; MV, PJCC and MIC collected data; MV analysed the data and drafted the manuscript; and PJCC, MIC, MLH and MDJ revised it. All authors approved the final version of the manuscript.

## Data archiving

Data with accompanying R code will be made available after manuscript acceptance.

## Supplementary Information

Table S1 and Annotated Registration.

## References

Agrawal AA (2000). Host-range evolution: Adaptation and trade-offs in fitness of mites on alternative hosts. Ecology 81: 500–508.

Agrawal AA, Conner JK, Rasmann S (2010). Tradeoffs and negative correlations in evolutionary ecology. In: Futuyma D, Levinton J, Eanes W (eds) Evolution since Darwin: the first 150 years, Sinauer, pp. 243–268.

Van Asch M, Julkunen-Tiito R, Visser ME (2010). Maternal effects in an insect herbivore as a mechanism to adapt to host plant phenology. FunctEcol 24: 1103–1109.

Berthouly A, Cassier A, Richner H(2008). Carotenoid-induced maternal effects interact with ectoparasite burden and brood size to shape the trade-off between growth and immunity in nestling great tits. Funct Ecol 22: 854–863.

Bilde T, Friberg U, Maklakov AA, Fry JD, Arnqvist G (2008). The genetic architecture of fitness in a seed beetle: Assessing the potential for indirect genetic benefits of female choice. BMC Evol Biol 8.

Bolker BM, Brooks ME, Clark CJ, Geange SW, Poulsen JR, Stevens MHH, et al. (2009). Generalized linear mixed models: a practical guide for ecology and evolution. Trends Ecol Evol 24: 127–35.

Cahenzli F, Erhardt A (2013). Transgenerational acclimatization in an herbivore-host plant relationship. Proc R Soc B Biol Sci 280.

Charmantier A, Garant D (2005). Environmental quality and evolutionary potential: Lessons from wild populations. Proc R Soc B BiolSci 272: 1415–1425.

Credland PF, Wright AW (1989). Factors affecting female fecundity in the cowpea seed beetle, Callosobruchus maculatus (Coleoptera: Bruchidae). J Stored Prod Res 25: 125–136.

Fox CW (1993). Maternal and genetic influences on egg size and larval performance in a seed beetle. Heredity 73: 509–517.

Fox CW, Bush ML, Roff DA, Wallin WG (2004). Evolutionary genetics of lifespan and mortality rates in two populations of the seed beetle, Callosobruchus maculatus. Heredity 92: 170–181.

Fox CW, Bush ML, Wallin WG (2003). Maternal age affects offspring lifespan of the seed beetle. Funct Ecol 17: 811–820.

Fox CW, Czesak ME (2000). Evolutionary ecology of progeny size in arthropods. Annu Rev Entomol 45: 341–369.

Fox CW, Dingle H (1994). Dietary mediation of maternal age effects on offspring performance in a seed beetle (Coleoptera: Bruchidae). Funct Ecol 8: 600–606.

Fox CW, Messina FJ (2018). Evolution of larval competitiveness and associated life-history traits in response to host shifts in a seed beetle. J Evol Biol 31: 302–313.

Fox CW, Savalli UM (1998). Inheritance of environmental variation in body size: Superparasitism of seeds affects progeny and grandprogeny body size via a nongenetic maternal effect. Evolution 52: 172.

Fox CW, Savalli UM (2000). Maternal effects mediate host expansion in a seed-feeding beetle. Ecology 81: 3–7.

Fox CW, Stillwell RC, Amarillo-Suárez AR, Czesak ME, Messina FJ (2004). Genetic architecture of population differences in oviposition behaviour of the seed beetle Callosobruchus maculatus. J Evol Biol 17: 1141–1151.

Fox CW, Thakar MS, Mousseau TA (1997). Egg size plasticity in a seed beetle: An adaptive maternal effect. Am Nat 149: 149–163.

Galloway LF (2005). Maternal effects provide phenotypic adaptation to local environmental conditions. New Phytol 166: 93–100.

Gompert Z, Messina FJ (2016). Genomic evidence that resource-based trade-offs limit host-range expansion in a seed beetle. Evolution 70: 1249–1264.

Gripenberg S, Mayhew PJ, Parnell M, Roslin T (2010). A meta-analysis of preference-performance relationships in phytophagous insects. Ecol Lett 13: 383–393.

Guntrip J, Sibly RM (1998). Phenotypic plasticity, genotype-by-environment interaction and the analysis of generalism and specialization in Callosobruchus maculatus. Heredity 81: 198–204.

Guntrip J, Sibly RM, Holloway GJ (1997). The effect of novel environment and sex on the additive genetic variation and covariation in and between emergence body weight and development period in the cowpea weevil, Callosobruchus maculatus (Coleoptera, Bruchidae). Heredity 78: 158–165.

Hadfield JD (2010). MCMC methods for multi-response generalized linear mixed models: The MCMCglmm R package. J Stat Softw 33.

Hansen TF, Pélabon C, Houle D (2011). Heritability is not evolvability. Evol Biol 38: 258–277.

Hill WG (2010). Understanding and using quantitative genetic variation. Philos Trans R Soc B Biol Sci 365: 73–85.

Hoffmann AA, Merilä J (1999). Heritable variation and evolution under favourable and unfavourable conditions. Trends Ecol Evol 14: 96–101.

Holloway GJ, Povey SR, Sibly RM (1990). The effect of new environment on adapted genetic architecture. Heredity 64: 323–330.

Houle D (1992). Comparing evolvability and variability of quantitative traits. Genetics 130: 195–204.

Iglesias-Carrasco M, Brookes S, Kruuk LEB, Head ML (2020). The effects of competition on fitness depend on the sex of both competitors. Ecol Evol 10: 9808–9826.

Iglesias-Carrasco M, Jennions MD, Zajitschek SRK, Head ML (2018). Are females in good condition better able to cope with costly males? Behav Ecol 29: 876–884.

Joshi A, Thompson JN (1995). Trade-offs and the evolution of host specialization. Evol Ecol 9: 82–92.

Kawecki TJ (1995). Expression of genetic and environmental variation for life history characters on the usual and novel hosts in Callosobruchus maculatus (Coleoptera: Bruchidae). Heredity 75: 70–76.

Kébé K, Alvarez N, Tuda M, Arnqvist G, Fox CW, Sembène M, et al. (2017). Global phylogeography of the insect pest Callosobruchus maculatus (Coleoptera: Bruchinae) relates to the history of its main host, Vigna unguiculata. J Biogeogr 44: 2515–2526.

Kruuk LEB (2004). Estimating genetic parameters in natural populations using the ‘animal model’. Philos Trans R Soc B Biol Sci 359: 873–890.

Kruuk LEB, Hadfield JD (2007). How to separate genetic and environmental causes of similarity between relatives. J Evol Biol 20: 1890–1903.

Leftwich PT, Nash WJ, Friend LA, Chapman T (2019). Contribution of maternal effects to dietary selection in Mediterranean fruit flies. Evolution 73: 278–292.

Lynch M, Walsh B (1998). Genetics and Analysis of Quantitative Traits. Sinauer.

McAdam AG, Garant D, Wilson AJ (2014) The effects of others’ genes: maternal and other indirect genetic effects. In:Charmantier A, Garant D, Kruuk LEB (eds) Quantitative Genetics in the Wild. Oxford University Press, pp. 81–104.

McGlothlin JW, Galloway LF (2014). The contribution of maternal effects to selection response: An empirical test of competing models. Evolution 68: 549–558.

Messina FJ (1993). Heritability and ‘evolvability’ of fitness components in Callosobruchus maculatus. Heredity (71: 623–629.

Messina FJ (2004a). Predictable modification of body size and competitive ability. Evolution 58: 2788–2797.

Messina FJ (2004b). How labile are the egg-laying preferences of seed beetles? Ecol Entomol 29: 318–326.

Messina FJ, Durham SL (2015). Loss of adaptation following reversion suggests trade-offs in host use by a seed beetle. J Evol Biol 28: 1882–1891.

Messina FJ, Fry JD (2003). Environment-dependent reversal of a life history trade-off in the seed beetle Callosobruchus maculatus. J Evol Biol 16: 501–509.

Messina FJ, Lish AM, Gompert Z (2018). Variable responses to novel hosts by populations of the seed beetle Callosobruchus maculatus (Coleoptera: Chrysomelidae: Bruchinae). Environ Entomol 47: 1194–1202.

Messina FJ, Morrey JL, Mendenhall M (2007). Why do host-deprived seed beetles ‘dump’ their eggs? Physiol Entomol 32: 259–267.

Messina FJ, Slade AF (1997). Inheritance of host-plant choice in the seed beetle Callosobruchus maculatus (Coleoptera: Bruchidae). Ann Entomol Soc Am 90: 848–855.

Mitchell R (1975). The evolution of oviposition tactics in the bean weevil, Callosobruchus maculatus (F.). Ecology 56: 696–702.

Moore MP, Whiteman HH, Martin RA (2019). A mother’s legacy: the strength of maternal effects in animal populations. Ecol Lett 22: 1620–1628.

Mousseau TA, Fox CW (1998). The adaptive significance of maternal effects. Trends Ecol Evol 13: 403–407.

Mousseau TA, Roff DA (1987). Natural selection and the heritability of fitness components. Heredity 59: 181–197.

Parichy DM, Kaplan RH (1992). Maternal effects on offspring growth and development depend on environmental quality in the frog Bombina orientalis. Oecologia 91: 579–586.

Paukku S, Kotiaho JS (2008). Female oviposition decisions and their impact on progeny life-history traits. J Insect Behav 21: 505–520.

Plaistow SJ, Benton TG (2009). The influence of context-dependent maternal effects on population dynamics: An experimental test. Philos Trans R Soc B Biol Sci 364: 1049–1058.

Price TN, Leonard A, Lancaster LT (2017). Warp-speed adaptation to novel hosts after 300 generations of enforced dietary specialisation in the seed beetle Callosobruchus maculatus (Coleoptera: Chrysomelidae: Bruchinae). Eur J Entomol 114: 257–266.

R Core Team (2020) R: A language and environment for statistical computing. v. 4.0.0. R Foundation for Statistical Computing, Vienna, Austria.

Räsänen K, Kruuk LEB (2007). Maternal effects and evolution at ecological time-scales. Funct Ecol 21: 408–421.

Roff DA, Wilson AJ (2014) Quantifying Genotype-by-Environment interactions in laboratory systems. In:Hunt J, Hosken D (eds) Genotype-by-Environment Interactions and Sexual Selection. John Wiley and Sons, Ltd., pp. 101–136.

Rossiter M (1996). Incidence and consequences of inherited environmental effects. Annu Rev Ecol Syst 27: 451–476.

Rossiter M (1998) The role of environmental variation in parental effects expression. In:Mousseau TA, Fox CW (eds) Maternal Effects as Addaptations. Oxford University Press, pp. 112–134.

Rowiński PK, Rogell B (2017). Environmental stress correlates with increases in both genetic and residual variances: A meta-analysis of animal studies. Evolution 71: 1339–1351.

Saltz JB, Bell AM, Flint J, Gomulkiewicz R, Hughes KA, Keagy J (2018). Why does the magnitude of genotype-by-environment interaction vary? EcolEvol 8: 6342–6353.

Scheirs J, Jordaens K, De Bruyn L (2005). Have genetic trade-offs in host use been overlooked in arthropods? Evol Ecol 19: 551–561.

Shama LNS, Strobel A, Mark FC, Wegner KM (2014). Transgenerational plasticity in marine sticklebacks: Maternal effects mediate impacts of a warming ocean. Funct Ecol 28: 1482–1493.

Spiegelhalter DJ, Best NG, Carlin BP, Van Der Linde A (2002). Bayesian measures of model complexity and fit. J R Stat Soc Ser B Stat Methodol 64: 583–616.

Stillwell RC, Wallin WG, Hitchcock LJ, Fox CW (2007). Phenotypic plasticity in a complex world: Interactive effects of food and temperature on fitness components of a seed beetle. Oecologia 153: 309–321.

Tucić N, Šešlija D (2007). Genetic architecture of differences in oviposition preference between ancestral and derived populations of the seed beetle Acanthoscelides obtectus. Heredity 98: 268–273.

Ueno H, Fujiyama N, Yao I, Sato Y, Katakura H (2003). Genetic architecture for normal and novel host-plant use in two local populations of the herbivorous ladybird beetle, Epilachna pustulosa. J Evol Biol 16: 883–895.

Vega-Trejo R, Head ML, Jennions MD, Kruuk LEB (2018). Maternal-by-environment but not genotype-by-environment interactions in a fish without parental care. Heredity 120: 154–167.

Via S, Lande R (1985). Genotype-environment interaction and the evolution of phenotypic plasticity. Evolution 39: 505–522.

de Villemereuil P, Schielzeth H, Nakagawa S, Morrissey MB (2016). General methods for evolutionary quantitative genetic inference from generalized mixed models. Genetics 204: 1281–1294.

Wasserman SS, Asami T (1985). The effect of maternal age upon fitness of progeny in the southern cowpea weevil, Callosobruchus maculatus. Oikos 45: 191–196.

Wilson AJ, Coltman DW, Pemberton JM, Overall ADJ, Byrne KA, Kruuk LEB (2005). Maternal genetic effects set the potential for evolution in a free-living vertebrate population. J Evol Biol 18: 405–414.

Wilson K, Hill L (1989). Factors affecting egg maturation in the bean weevil Callosobruchus maculatus. Physiol Entomol 14: 115–126.

Wilson AJ, Réale D, Clements MN, Morrissey MB, Postma E, Walling CA, et al. (2010). An ecologist’s guide to the animal model. J Anim Ecol79: 13–26.

Wolf JB, Wade MJ (2009). What are maternal effects (and what are they not)? Philos Trans R Soc London B 364: 1107–1115.

Wood CW, Wice EW, del Sol J, Paul S, Sanderson BJ, Brodie ED (2018). Constraints imposed by a natural landscape override offspring fitness effects to shape oviposition decisions in wild forked fungus beetles. Am Nat 191: 524–538.

