## Supplementary Information for "The role of maternal effects on offspring performance in familiar and novel environments"

Supplementary Information for Vrtílek et al. - **The role of maternal effects on offspring performance in familiar and novel environments**

Table S1. Summary of offspring trait averages and the sample size per host type used in the study.

Annotated registration of '*Interaction between maternal and environmental effects in a seed beetle*', the original version can be found at the link: https://osf.io/ft7eq/?view_only=0bab0a33bb4246adb64c919601a72757

**Table S1.** Summary of offspring trait averages (± standard deviation) and the sample size per host type used in the study.

| Trait | Sex | Original host (cowpea) | N (% of total) | Novel host (mung bean) | N |
|---|---|---|---|---|---|
| Larval survival (probability) | - | 0.889 | 1 727 (55 %) | 0.930 | 1 431 |
| Duration of larval development (days) | Females (daughters) | 32.025 ± 1.355 | 734 (52 %) | 30.770 ± 1.009 | 686 |
| | Males (sons) | 31.526 ± 1.481 | 741 (54 %) | 30.456 ± 1.102 | 615 |
| Body mass (mg) | Females (daughters) | 5.826 ± 0.707 | 743 (52 %) | 6.515 ± 0.615 | 688 |
| | Males (sons) | 3.527 ± 0.564 | 761 (54 %) | 3.885 ± 0.459 | 617 |
| Adult lifespan (days) | Females (daughters) | 22.926 ± 6.368 | 744 (52 %) | 26.189 ± 6.129 | 681 |
| | Males (sons) | 16.913 ± 5.367 | 786 (57 %) | 19.366 ± 5.164 | 621 |
| Relative host preference (eggs on cowpea:total eggs ratio) | Females (daughters) | 0.681 ± 0.187 | 364 (55 %) | 0.517 ± 0.178 | 301 |

### Annotated registration of 'Interaction between maternal and environmental effects in a seed beetle'

[November 19, 2020 edits in RED]

Original version: https://osf.io/ft7eq/?view_only=0bab0a33bb4246adb64c919601a72757

**Study stage:** post-analysis phase (data analysed and interpreted)

**Main questions**

Is there variation among females in the relative success of their offspring in two host environments?

Is relative offspring success on each host predicted by a mother preference for that host?

**Hypotheses** (order and formulations have changed to better reflect focus of the study and the different analytical approach that we used)

1) Maternal effect – There is a strong maternal effect on offspring performance in seed beetle

Prediction: Among-female (dam effect) variation in offspring performance will be greater than variation among families (sire effect)

1) There are strong maternal effects on offspring life-history traits. This led us to predict that a considerable component of offspring phenotypic variation, beyond that due to additive genetic effects, would be due to offspring having different mothers.

2) M×E interaction – The extent to which the original host is superior to the new host for rearing offspring differs among individual dams

Prediction: Variation among dams in offspring performance (see Hypothesis 1) will differ between novel and original host environment resulting in a significant M×E interaction

2) Maternal effects are environment-specific (M×E). Specifically, we predicted that the maternal effects variance would differ between offspring host environments, and/or that the rank of offspring from the same mother will change between the two host types decreasing the cross-environmental maternal correlation.

3) Host effect – A novel host environment will reduce offspring performance

Prediction: Offspring from the novel host will have lower egg-to-adult survival, lower adult body mass, longer larval development, and shorter adult lifespan compared to the original host

3) Novel host type will be more challenging for offspring. Hence, offspring developing in the novel host would suffer reduced performance compared to those developing in

the original host type (i.e. lower larval survival, longer larval development, lower body mass at emergence, and shorter adult lifespan).

4) Adaptiveness of host preference – Offspring performance on different hosts will be positively correlated with dam host preference

Prediction: The extent to which mothers prefer to lay eggs on the original versus the novel environment will be positively correlated with offspring performance on the original host, but negatively or uncorrelated to the performance on the novel host.

4) Maternal host preference will predict offspring performance. We predicted that offspring would perform better on the host type preferred by their mother when she laid her eggs.

**Sample size**

Our sample size (the number of $F_1$ offspring) was determined a priori based on Lynch & Walsh (1998)[1]. We mated 356 dams and 89 sires (1 sire per 4 dams) and aimed for 10 offspring per dam per host species. Our final sample size is lower, however, mainly due to unsuccessful mating and no eggs laid in preference trial and/or on one of the host types.

Overall, we collected 3494 seeds with a single egg laid, out of which 2935 came from mothers that had laid at least one egg both during the preference trial and also on both host types. We then recorded data for 2639 emerged offspring (1306 males and 1333 females). For the analysis of heritability, we quantified host preference for 622 female offspring that also had information on dam host preference.

The data are not yet examined and therefore, no data has been removed based on the distribution and no data-transformation has been applied. If we identify any clear typo-outliers (e.g. 1234 mg recorded instead of 1.234 mg for body mass), we will correct/remove them. Outliers should be properly handled by linear mixed models as these models weigh the value of the effect by the sample size across hierarchical levels. We will remove extreme values further than 3 standard deviations from the mean. Distribution of residuals will be examined with diagnostic plots of linear mixed models and if the residuals are non-normally distributed, generalized linear mixed models will be applied instead. Some of the variables (e.g. survival – died/emerged) are obviously non-Gaussian and we will apply generalized linear mixed model directly.

[1]Lynch M & Walsh B (1998) *Genetics and Analysis of Quantitative Traits*. Sinauer Associates. Sunderland, MA, USA. pp 980.

(The sample sizes differ in the final analyses as we decided to analyse the two sexes separately and not to remove offspring of dams who had offspring from one host type only. We still excluded offspring with extreme values (>3 standard deviations from the mean) and offspring of dams without recorded host preference (N = 285). Sample size therefore varies among individual life-history traits: 'larval survival' (dead or emerged; N = 3146), 'duration of larval development' (number of days between oviposition and offspring emergence; N = 1 420 daughters and 1 384 sons), 'body mass' (weight at emergence in mg; N = 1 431 daughters and 1 399 sons) and 'adult lifespan' (number of days between offspring emergence and death; N = 1 425 daughters and 1 389 sons). In addition, we measured the host egg laying preferences of 665 daughters.)

**Variables measured**

Response variables:

offspring performance

- offspring egg-to-emergence survival (died/emerged)
- offspring body mass after emergence from bean (weight to 0.001mg)
- offspring larval development duration (number of days between egg laying and emergence)
- offspring adult lifespan (number of days between emergence and death)

  here, the sample size is 488 for females; we excluded females that were mated (to measure their host preference), because copulation was previously shown to affect lifespan in seed beetles

(We did not remove mated females in the analysis of adult lifespan, so the sample size for daughters was 1 425.)

offspring host preference

- relative preference for the original host type (number of eggs laid on the original/total eggs laid)

Explanatory variables and covariates used in the Main analyses:

Continuous variables will be scaled to zero mean and standard deviation of 1 to facilitate model computation

- host (treatment effect = original/novel)
- offspring sex (male/female)
- dam mating order (1-4, ordinal scale)
- seed mass (weight of the seed after egg has been laid, weight to 0.001mg)

  we will standardize the values per each host to zero host mean and 1 SD for host variance
- dam relative preference for the original host type (number of eggs laid on the original/total eggs laid)

(We added continuous fixed effect 'day mated' with values 1-6 in the 'minimal' and 'full' models to account for the experimental block effect originally fitted in the random part of the model. This was due to computing problems when the random effects part of the mixed-effects model became overly complex and the low number of levels.)

Additional explanatory variables and covariates that might be used in Exploratory analyses:
(these are not used in the final analysis)

- offspring body mass
- offspring larval development duration
- dam body mass (weight to 0.001mg)

- sire body mass (weight to 0.001mg)
- sire and dam adult lifespan (number of days between egg laying and emergence)

Random effects:

- sire ID, dam ID
- BLOCK (date parents were mated)

  optional – BLOCK will be dropped if model fails to converge

(BLOCK was not included, but fitted as a continuous fixed effect 'day mated'. See above.)

**Main analyses** (in R software using packages *lme4*, *nlme*)

We aim at partitioning phenotypic variance among the effect of genetic additive variance (sire effect) and maternal and dominance effects (dam effect) and their interaction with environment (host). We will therefore initially omit parent and offspring properties that might explain variation in focal offspring performance traits, and only include covariates that vary due to the experimental design (e.g. whether the dam was a male's first or last mate). We will build a set of models with fixed effects held constant but with different structure of random effects reflecting the hierarchical character of our half-sib/full-sib breeding design. Our goal is to evaluate the 'full' model (containing both genotype-by-environment and maternal-by-environment interactions) compared to models with reduced random effect structure using similar approach to Vega-Trejo et al. (2018)[2].

To eventually test for (1) maternal effect (M), (2) host effect (E), and (3) M×E interaction, we will apply model selection based on Akaike information criterion (AIC) to establish the best random-effects structure (i.e. lowest AIC). We will consider a difference of 2 units of AIC as biological significant. Then, we will assess the effect of fixed terms in the top-selected model based on their confidence intervals and inferential statistics ($\alpha$ = 0.05). If the SEX:HOST interaction term is significant, we will analyse sexes separately.

To test for (4) the effect of dam host preference on her offspring performance, we will include 'HOST×dam relative host preference' interaction instead of just 'HOST' alone. We will then perform the model selection as described above for (1-3).

[2]Vega-Trejo R, Head ML, Jennions MD, Kruuk LEB (2018) Maternal-by-environment but not genotype-by-environment interactions in a fish without parental care. *Heredity* **120**: 154–167.

| **Model** | **Structure** |
|---|---|
| $M_{full}$ | offspring performance ~ HOST + covariates + (HOST\|ID♀) + (HOST\|ID♂) |
| $M_{G\times E\ only}$ | offspring performance ~ HOST + covariates + (1\|ID♀) + (HOST\|ID♂) |
| $M_{M\times E\ only}$ | offspring performance ~ HOST + covariates + (HOST\|ID♀) + (1\|ID♂) |
| $M_{M+G\ only}$ | offspring performance ~ HOST + covariates + (1\|ID♀) + (1\|ID♂) |
| $M_{G\ only}$ | offspring performance ~ HOST + covariates + (1\|ID♂) |
| $M_{null}$ | offspring performance ~ HOST + covariates |

offspring performance: egg-to-emergence survival, body mass, larval development duration, adult lifespan

covariates: SEX, seed mass and their interactions with HOST (SEX:HOST, seed mass:HOST), dam mating order

Terms in parentheses denote the random effects as: (random slope|random intercept); '(1|BLOCK)' will be included as random intercept if it does not prohibit model convergence due to overparametrization. If the model does not converge block will be dropped from the model.

(We did not follow the original approach and fit MCMC animal model instead (using R package MCMCglmm, ver. 2.29[3]). We used random effects structure based on a pedigree of beetles in our experiment. To model maternal effects above additive genetic effects, we include dam ID. We define a set of candidate models that we compare using Deviance Information Criterion (DIC) that is similar to AIC.

The simplest model 'G' included only additive genetic effects specified by random effect of 'animal'. We then included also dam identity to estimate maternal effects in addition the additive genetic effects ('G+M' model). To estimate each variance component separately per host type we specified random-effect interaction (Hadfield, 2010) with host type 'idh(HOST):animal' for 'Gsep', or with 'idh(HOST):dam' for 'Msep' models. This is similar to fitting two separate host-specific models. We further fitted models that also estimated cross-environmental covariance ('Gcov' or 'Mcov'). We used 'unstructured' variance-covariance matrix, as e.g. 'us(HOST):animal' (Hadfield, 2010), so we also obtained the corresponding covariance to compute the cross-environmental correlations – the proxies for Genotype-by-Environment (G×E) or Maternal-by-Environment (M×E) interaction (Lynch & Walsh 1998). In the models estimating separate variances per host type, we always specified host-specific residual variance (as 'rcov=~idh(HOST):units') (Hadfield, 2010).

[3]Hadfield JD (2010) MCMC methods for multi-response generalized linear mixed models: The MCMCglmm R package. *J Stat Softw* **33**.)

Model specification for model comparison:

| **Model** | **Structure** |
|---|---|
| G | offspring response ~ HOST + covariates + animal |
| G+M | offspring response ~ HOST + covariates + animal + dam |
| $G_{sep}$ | offspring response ~ HOST + covariates + idh(HOST):animal |
| $G_{cov}$ | offspring response ~ HOST + covariates + us(HOST): animal |
| $G_{sep}$+$M_{sep}$ | offspring response ~ HOST + covariates + idh(HOST):animal + idh(HOST):dam |
| $G_{cov}$+$M_{sep}$ | offspring response ~ HOST + covariates + us(HOST): animal + idh(HOST):dam |
| $G_{cov}$+$M_{cov}$ | offspring response ~ HOST + covariates + us(HOST): animal + us(HOST):dam |

offspring response: egg-to-emergence survival, body mass, larval development duration, adult lifespan, host preference; covariates: bean mass, dam mating order, day mated

We use the random-effects structure of the best candidate model to fit minimal and full model. The 'minimal model' contains only terms coming from the experimental design and allows us to estimate terms of phenotypic variance partitioning. In the 'full model', we test the effect of additional explanatory variables on offspring performance.

Interpretation of potential results:

Hyp.1: Strong maternal effect will be indicated by higher variance proportion explained by ID♀ compared to ID♂ in the $M_{M+G\ only}$ model. We will test this directly using likelihood-ratio test (LRT) comparing $M_{M+G\ only}$ vs. $M_{G\ only}$.

(Strong maternal effects would be indicated by models containing dam-related variance showing low DIC values. We would consider DIC difference between two models of 2 as 'significant'.)

Hyp.2: We will consider M×E interaction important if it is the top-selected model based on the AIC comparison. We will test the importance directly using LRT comparing $M_{G\times E\ only}$ vs. $M_{full}$.

(We will consider M×E interaction important if models with $M_{sep}$ or $M_{cov}$ are in the top-selected models based on the DIC comparison, or the cross-environmental maternal effect correlation is lower than 0.999 based on the credible intervals from posterior model distribution. The cross-environmental maternal effect correlation is calculated as $r_M = \frac{COV_{M\ (original-nove\ )}}{\sqrt{V_{M\ (original)} V_{M\ (novel)}}}$.)

Hyp.3: The host effect will be tested by the main HOST term in the fixed part of the model. If significant, then one host is better for offspring performance than the other.

Hyp.4: We can conclude that there is an effect of mother preference on offspring performance if the 'HOST×dam relative host preference' interaction is significant. We expect a positive effect of 'dam relative host preference' on offspring performance from the original host – the more the original host is preferred, the better the relative performance of offspring on that host.

Estimate of heritability:

We will quantify heritability of both offspring performance traits and host preference. If G×E comes out as significant, we will estimate heritability separately for each host, otherwise we will pool the offspring together. We will only specify random effects of dam and sire to decompose the variance among families (paternal half-sibs) and full-sibs (also using block if possible). For example, for host preference:

daughter relative host preference ~ seed mass + HOST + seed mass:HOST + dam mating order + (1|ID♀) + (1|ID♂:ID♀) + (1|BLOCK)

$V_A$ = 4x variation among families (sires, (1|ID♂:ID♀))

$h^2 = V_A/V_{TOT}$

(We estimated heritability in the two host types separately based on formula $h^2 = \frac{V_A}{V_{total}} = \frac{V_{animal}}{(V_{animal} + V_{dam} + V_{residual})}$. The Genotype-by-Environment interaction was tested in the final analysis in the form of correlation between genetic effects across the two host types. We fitted covariance of additive genetic effects at the two host types and calculated $r_G = \frac{COV_{animal\ (original-nove\ )}}{\sqrt{V_{animal\ (original)} V_{animal\ (novel)}}}$. We also provide evolvability (standardized additive genetic variance across trait mean) as $CV_A = \frac{\sqrt{V_A}}{\mu} = \frac{\sqrt{V_{animal}}}{\mu}$, where μ is trait mean.)

Effect of dam condition on offspring performance: (not tested in the final analysis)

We will further test whether dam body mass affected the offspring performance positively and whether this effect is stable in the two host environments. We will fit 'dam body mass×HOST interaction' as a fixed term in the above described model selection procedure.

Effect of offspring development duration on body mass: (not tested in the final analysis)

In addition, we expect that offspring body mass and offspring development duration will be negatively correlated. We plan to test this across the two hosts by specifying 'offspring development duration×HOST' interaction in model of offspring body mass as response variable and use the model selection procedure as above.

**Secondary (exploratory) analyses** (not conducted)

Here is a list of four parental properties that might covary with the offspring responses but were not included as fixed factors in the main analysis of phenotypic variance decomposition.

- sire body mass (weight to 0.001mg)
- sire adult lifespan (number of days between egg laying and emergence)
- dam adult lifespan (number of days between egg laying and emergence)

We will use these traits as covariates in exploratory analyses to test whether they predict variation in the three focal offspring performance traits of:

- offspring body mass after emergence from bean (weight to 0.001mg)
- offspring larval development duration (number of days between egg laying and emergence)
- offspring adult lifespan (number of days between emergence and death)
